# GTDB-based packages enhance GraftM performance

**DOI:** 10.1101/2023.10.03.560708

**Authors:** Aleksei A. Korzhenkov

## Abstract

In this short note I present a way to enhance GraftM performance and incorporate novel genome-based prokaryotic taxonomy to sequencing data classification.

## Introduction

GraftM performs taxonomic and functional classification of metagenomic reads using packages for specific genes (Boyd, J.A., Woodcroft, B.J., Tyson G.W., 2018). According to Dimensions (https://badge.dimensions.ai/details/id/pub.1101437024) GraftM is still actual software having 34% of 116 citations in the past two years. 61 packages for taxonomic classifications are available in the specific GraftM gpkgs repository (https://github.com/geronimp/graftm_gpkgs/), but most of the packages were uploaded six years ago and a small part four years ago. Fortunately, GraftM provides users an ability to construct their own packages, supplying the program with reference sequences in FASTA format, taxonomy as CSV file, and optional HMM profile for gene searching.

## Methods

Amino acid sequences of Ribosomal protein L10 (PF00466) and Ribosomal protein S8 (PF00410) were taken from representative genomes of both archaea and bacteria from the GTDB R207 database (Parks, D.H., et al. 2021). Redundant sequences were filtered using CD-HIT (Fu, L., et al., 2012) with identity threshold set to 1 and all other settings set to default. Taxonomy file was combined from separate bacterial and archaeal genome-based taxonomies. New GraftM packages were built according to the software manual.

Resulting packages and reference package for ribosomal protein L10 from the GraftM gpkgs repository were tested on publicly available data of shotgun metagenome sequencing of microbial mats from alkaline hot spring (NCBI SRA accession SRR6429753) analyzed earlier (Korzhenkov, A.A., et al., 2018). Sequencing reads were trimmed by quality (q=20), adapters and poly-G regions were removed and short reads (ml=33) were discarded using BBDuk (https://sourceforge.net/projects/bbmap/).

## Results

57 756 sequences for L10 and 57 667 for S8 genes from archaeal and bacterial representative genomes of GTDB were taken to remove redundancy leaving 52 844 and 47 578 sequences for L10 and S8 genes respectively. In comparison, the original L10 package includes 6 215 sequences. Larger amount of data used to construct new packages leads to growth of reference phylogenetic tree and increased duration of tree insertion step of GraftM pipeline carried by pplacer. However it takes only four minutes or less (Table 1) to classify the dataset with the new L10 package on a modern PC (four threads of AMD 3900 CPU were used).

**Table 1.** Control dataset classification results

|  | **L10 (original)** | **L10 GTDB** | **S8 GTDB** |
| --- | --- | --- | --- |
| **reads placed in tree** | 174 | 175 | 197 |
| **reads classified to genus** | 12 | 118 | 107 |
| **genus count** | 3 | 8 | 12 |
| **reads classified to family** | 47 | 48 | 59 |
| **family count** | 7 | 13 | 21 |
| **reads classified to family or genus, %** | 33.9 | 94.9 | 84.3 |
| **reads classified to order** | 24 | 0 | 12 |
| **order count** | 9 | 11 | 20 |
| **runtime, seconds** | 25.38 | 242.6 | 208.47 |

Sequences of metagenome assembled genomes (MAGs) included in GTDB help classify uncultured prokaryotes, which are a significant part of microbial communities. The sensitivity of new packages is comparable with the old one. New packages show better resolution on genus and family ranks: share of reads classified to rank of genus or family increased from 33.9% to 84.3 (S8) and 94.9% (L10) (Table 1). GraftM GTDB-based packages are freely available in the GitHub repository https://github.com/laxeye/graftm-add-gpkg.

**Figure 1.**
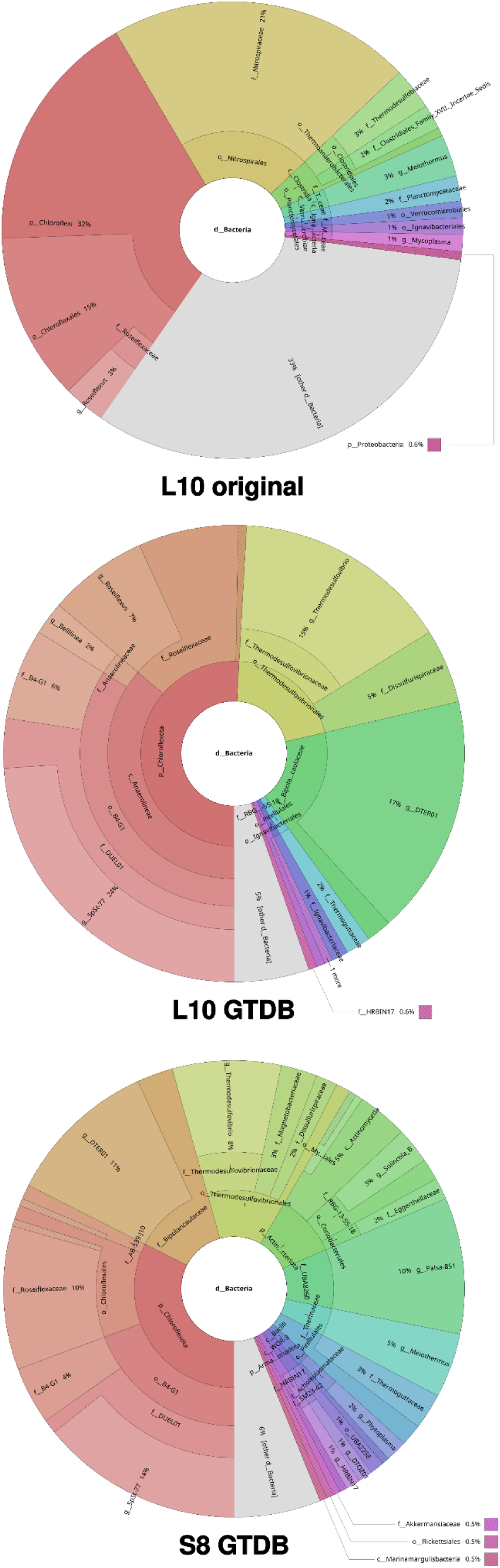
Taxonomy profiles of SRR6429753 data calculated with original L10 (top) and new L10 (middle) and S8 (bottom) GTDB packages. Only bacterial data shown.

